# Endosymbiotic bacteria are prevalent and diverse in agricultural spiders

**DOI:** 10.1101/586750

**Authors:** Jennifer A. White, Alexander Styer, Laura C. Rosenwald, Meghan M. Curry, Kelton D. Welch, Kacie J. Athey, Eric G. Chapman

**Author notes:** Correspondence to: Jen White, Dept. of Entomology, Univ. of Kentucky, Lexington, KY 40546, ORCID: 0000-0003-1276-561X. The nucleotide sequence data reported are available in the DDBJ/EMBL/GenBank database under accession numbers MK513512-19, MK521215-57, MK529759-77, MK627518-20, MK631910-40, MK632848-60 and the NCBI SRA under PRJNA526944.

## Abstract

Maternally inherited bacterial endosymbionts are common in arthropods, but their distribution and prevalence is poorly characterized in many host taxa. For example, spiders (Araneae) have received little attention, but initial surveys suggest that vertically transmitted symbionts may be common. Here, we characterized endosymbiont infection in a community of agricultural spiders. Using a combination of diagnostic PCR and high-throughput sequencing of the bacterial microbiome, we evaluated symbiont infection in 267 individual spiders representing 14 species in 3 families. We found 27 Operational Taxonomic Units (OTUs) that are likely endosymbiotic, including several strains of *Wolbachia*, *Rickettsia* and *Cardinium*, all of which are vertically transmitted and frequently associated with reproductive manipulation of arthropod hosts. Seventy percent of spider species had individuals that tested positive for one or more endosymbiotic OTUs, and specimens frequently contained multiple symbiotic strain types. The most symbiont-rich species, *Idionella rugosa*, had eight endosymbiotic OTUs, with as many as five present in the same specimen. Individual specimens within infected spider species had a variety of symbiotypes, differing from one another in the presence or absence of symbiotic strains. Our sample included both starved and unstarved specimens, and dominant bacterial OTUs were consistent per host species, regardless of feeding status. We conclude that spiders contain a remarkably diverse symbiotic microbiota. Spiders would be an informative group for investigating endosymbiont population dynamics in time and space, and unstarved specimens collected for other purposes (e.g., food web studies) could be used, with caution, for such investigations.

## Introduction

The majority of terrestrial arthropod species are infected by inherited bacterial symbionts. The most common of such bacteria, *Wolbachia*, is estimated to infect approximately half of arthropod species, albeit often at a low frequency of infected individuals within a given species [1–3]. These bacteria are primarily transmitted vertically, but horizontal transfers among host taxa occasionally occur through a variety of ecological mechanisms such as shared host substrates or shared enemies [4–8]. Following successful establishment in a new host lineage, the endosymbiont then promotes the continued existence of that infected lineage, either by providing some sort of fitness benefit to the host [9,10], or by manipulating host reproduction such that infected mothers produce more daughters than uninfected mothers [11]. Over generations, the result is an infection that can spread through the host population via selection, rather than contagion.

It is inevitable that this process of infection and spread sometimes occurs in the context of other vertically-transmitted endosymbionts, either through co-infection of the same host individual, or through interactions of differentially-infected individuals in the host population. As molecular techniques for characterizing bacterial communities continue to improve, evidence is accumulating that symbiont diversity within and among hosts likely greater than previously appreciated [12–14]. While the dynamics of interacting symbionts have been considered theoretically in various contexts [e.g., 15,16], empirical work has been quite limited, perhaps because the dynamic phase of symbiont spread in host populations can be temporally transient [10,17]. Among insects, there are several examples of species with notably diverse endosymbiont complements (e.g., *Acyrthosiphum pisum*, *Bemisia tabaci*), although this diversity is often strongly structured among geographic populations or distinct biotypes [18,19]. Among spiders, much less is known about the dynamics and diversity of endosymbionts than in insects. However, the small body of literature on spider symbionts suggests that that may have a higher frequency of infection and diversity of symbionts than is typical in insects [20–27], and may contain novel symbiotic microbial taxa [13,27]. Thus, spiders may be of particular utility for exploring interactions among symbionts.

Our primary objective in this paper was to generate baseline data regarding endosymbiont diversity and variation among spiders. We used a combination of diagnostic PCR and high-throughput sequencing to characterize the endosymbiont community of common agricultural spiders, primarily within the family Linyphiidae. As a secondary objective, we contrasted the microbial community of starved versus unstarved spider specimens. As predators, spiders have the potential to give false positives for endosymbiont infection, due to consumption of symbiont-infected-prey. Thus, starved specimens with minimal gut contents are the most reliable indicators of genuine infection. However, unstarved specimens that were used in ecological or phylogenetic studies, e.g., [28,29], could be a useful repository of samples for subsequent studies of endosymbionts. Here we evaluate the potential utility of such specimens for symbiont characterization.

## Materials and Methods

Spider specimens originated from one of two groups [Online Resource 1]. Spiders in the first group were field-collected into individual microcentrifuge tubes, returned to the laboratory, and held in isolation for four or five days before being preserved in 95% ethanol. These specimens were designated as “starved” and were the least likely to include false positives due to symbiotic associates of undigested prey. The second group of “unstarved” spiders had been collected as part of foodweb experiments [K. Athey unpublished data, 29], and had been immediately frozen in the field with gut contents intact. All spiders originated from Fayette County, Kentucky, USA, from alfalfa, wheat, melon, or field margins at the University of Kentucky Horticultural Research Farm (37°58’39”N, 84°32’30”W) or Spindletop Research Farm (38°7’29”N, 84°30’39”W). A total of 267 specimens, representing 14 species in 3 families, were included in the study (Table 1).

**Table 1.** Results from diagnostic screen for three bacterial endosymbiotic genera in spiders

|  | Total<br>sampled | Infected with |  |  |  |
| --- | --- | --- | --- | --- | --- |
|  |  | *Cardinium* | *Wolbachia* | *Rickettsia* | Multiple<sup>a</sup> |
| Linyphiidae |  |  |  |  |  |
| *Agyneta semipallida* (Chamberlin & Ivie 1944) | 1 | 0 | 0 | 0 | 0 |
| *Agyneta serrata* (Emerton 1909) | 16 | 0 | 0 | 0 | 0 |
| *Agyneta unimaculata* (Banks 1892) | 5 | 1 | 0 | 2 | 0 |
| *Erigone autumnalis* Emerton 1882 | 21 | 0 | 2 | 0 | 0 |
| *Florinda coccinea* (Hentz 1850) | 4 | 0 | 0 | 0 | 0 |
| *Idionella rugosa* (Crosby 1905) | 75 | 75 | 63 | 20 | 65 |
| *Islandiana flaveola* (Banks 1892) | 2 | 0 | 0 | 0 | 0 |
| *Mermessus* sp. 1 | 25 | 0 | 24 | 2 | 2 |
| *Mermessus tridentatus* (Emerton 1882) | 1 | 0 | 1 | 1 | 1 |
| *Mermessus trilobatus* (Emerton 1882) | 66 | 1 | 13 | 45 | 12 |
| *Tennesseellum formica* Emerton 1882 | 20 | 0 | 1 | 0 | 0 |
| *Walckenaeria microspiralis* Milidge 1983 | 1 | 1 | 1 | 0 | 1 |
| Tetragnathidae |  |  |  |  |  |
| *Glenognatha foxi* (McCook 1894) | 20 | 0 | 6 | 0 | 0 |
| Oxyopidae |  |  |  |  |  |
| *Oxyopes salticus* Hentz 1845 | 10 | 0 | 7 | 0 | 0 |

<sup>a</sup>Individual spider specimens that tested positive for more than one symbiont genus

We surface-sterilized each specimen with a series of bleach and ethanol rinses [26], before extracting DNA from the abdomen using DNEasy Blood and Tissue extraction kits (Qiagen, Germantown, MD) according to manufacturer’s instructions. Extraction quality for each sample was verified by PCR amplification of a segment of the COI barcoding gene [Online Resource 2]. For group 1, this segment was sequenced for all spiders and compared to NCBI and BOLD databases to validate species identity. A subset of specimens from group 2 were also sequenced to verify previous morphological identifications [29].

We diagnostically screened all samples for three bacterial genera known for endosymbiotic associations with arthropods. We used previously published primers specific to *Cardinium* (Class Bacteroidetes, Order Bacteroidales, Family Bacteroidaceae), *Wolbachia* (Class Alphaproteobacteria, Order Rickettsiales, Family Anaplasmataceae), and *Rickettsia* (Class Alphaproteobacteria, Order Rickettsiales, Family Rickettsiaceae) [Online Resource 2]. All reactions were run in 10µl volume, with products electrophoresed and visualized on 1% agarose gels stained with Gel Red (Biotium) alongside known positive and negative (reagents-only) controls. Samples with initial negative diagnoses for a given bacterial genus were retested (with a different primer set, if possible) before being categorized as uninfected. For a subset of the samples with positive evidence of infection, we repeated the PCR at a 20µl volume and purified the PCR product with either GenCatch PCR Cleanup or Gel Extraction Kits (Epoch Life Sciences, Missouri City, TX) according to manufacturer’s instructions. All products were then submitted for Sanger sequencing at the University of Kentucky genomics core facility or an offsite sequencing facility (GENEWIZ, South Plainfield, NJ). Resulting sequences were compared to the NCBI nucleotide database using the megablast algorithm, and specimens returning a 97% or higher match to the expected bacterial genus were scored as positive. For several *Wobachia* strains, we additionally sequenced five genes used for MLST strain typing, using primers and procedures as described at https://pubmlst.org/wolbachia/ [30].

To investigate whether additional symbionts were present in these specimens, as well as to validate diagnostic results, we also profiled the microbiomes of a subset of 110 specimens using high-throughput sequencing of the bacterial community. We amplified the V4 region of bacterial 16S rRNA from a subset of samples that included both starved and unstarved specimens [Online Resource 1]. Each sample was individually tagged with a unique combination of indexed forward and reverse primers [31] that were multiplexed into one of three libraries that were each sequenced on Illumina Miseq 2500 instruments. Each library also included specimens from other projects that are not reported here, and received a PhiX spike to increase sequence heterogeneity among the amplified sequences. Two of the sequencing runs were conducted at the University of Kentucky genomics core facility, and the third at the B-CELL sequencing facility (Bluegrass Community & Technical College, Lexington, Kentucky). Several samples were included in more than one sequencing run, as a quality control.

Sequences from each run were demultiplexed, trimmed and quality filtered within BaseSpace (Illumina, basespace.illumina.com), then imported into qiime2 (v2017.11, https://qiime2.org; [32]) using a manifest. We conducted additional quality control using deblur [33], implemented in qiime2 using default parameters and a trim length of 251 bases. This procedure allowed us to distinguish among strain types (Operational Taxonomic Units, or OTUs) even if they only differed by one basepair over the sequenced length [34]. Resulting unique sequences were taxonomically classified using a naïve Bayes classifier that was trained on the 515F/806R V4 region of the Greengenes 13_8 99% OTUs reference database [35]. We additionally queried high prevalence sequences (>0.1% of total reads, or >1% of any particular sample) against the NCBI nt database using the megablast algorithm, to identify symbiotic taxa that may not have been included in the reference database. When the high-throughput data indicated the presence of other bacterial genera that are known to be vertically transmitted endosymbionts in arthropod hosts, (*Rickettsiella* and *Spiroplasma*) we conducted secondary diagnostic tests within several spider species to investigate frequency of infection among specimens that had not been included in the high-throughput dataset [Online Resource 2].

Individual specimens were scored for the presence of symbiotic taxa based on the combination of diagnostic, Sanger and high-throughput sequencing data. For spider species in which sequencing results consistently and reliably validated diagnostic PCR results, we also accepted the positive diagnoses of unsequenced specimens. When diagnostic positives could not be validated by either sequencing methodology, we reclassified them as negative. For high-throughput data, each sequencing run included samples that shared either forward and reverse indices with other samples in the same run. In the sequencing process, a detectable number of reads undergo index swapping, and get misallocated to the wrong sample [36]. To combat the potential for false positives, we took several precautions. First, we excluded all high-throughput samples with less than 1000 reads from further analysis (15/110 specimens). Second, for each sample, we only evaluated OTUs that constituted greater than 1% of reads, and totaled more than 100 reads for that sample. Third, for the OTUs that were most abundant overall in a given run (>1% of total reads), we used the number of reads found in several blank and control samples as a benchmark, and only considered a specimen potentially positive for that OTU if it had more than 2× the maximum number of reads in the negative controls. This conservative approach resulted in good correspondence with diagnostic and Sanger sequencing results, but potentially resulted in some false negatives in genuinely infected specimens, particularly for OTUs that had high representation in other specimens in the sequencing run.

For two spider species (*Idionella rugosa* and *Glenognatha foxi*) we had enough specimens in both starved and unstarved categories to merit a comparison between feeding statuses, although it should be noted that this was not a randomly imposed treatment, and the data should be interpreted with caution. Nevertheless, as an exploratory measure we calculated bacterial community richness in starved and unstarved specimens in each of these species using Shannon’s Index and observed OTUs for samples rarified to a uniform depth of 3000 reads in qiime2’s core phylogenetics package, and compared values between feeding status categories using nonparametric Kruskal-Wallis tests. Bacterial community composition was compared between categories via PERMANOVA of both weighted and unweighted UniFrac values [37].

## Results

Endosymbionts were very common in the agricultural spiders that we screened. Diagnostic screening results are presented in Table 1. Most spider species (10/14 species = 71%) had at least one specimen that tested positive for common endosymbiont genera, and half (7/14) of the species had two or more symbiont genera represented. Five species had specimens that simultaneously tested positive for symbionts of multiple genera, likely representing co-infections. For most species, at least some of the positive diagnoses resulted from specimens that were starved for several days prior to extraction [Online Resource 1], and thus likely represent true infections, rather than false positives due to symbiont-infected prey in the gut. Further comparison of starved versus unstarved specimens is included below.

When we used sequencing data to distinguish among symbiont strains, we found that multiple strains of each endosymbiotic genus were represented (Table 2, [Online Resources 1,3]), often within the same spider specimen. Fig. 1 depicts the symbiont profiles of individual specimens from three of the most diversely-infected spider species, as determined by high-throughput sequencing. In *I. rugosa*, we detected eight symbiont strains, up to five of which co-occurred within individual specimens (Fig. 1a). In addition to two strains each of *Cardinium*, *Wolbachia*, and *Rickettsia*, we also detected strains of *Rickettsiella* (Class Gammaproteobacteria, Order Legionellales, Family Coxiellaceae) and *Rhabdochlamydia* (Class Chlamydiae, Order Chlamydiales, Family Rhabdochlamydiaceae), both of which have been associated with symbiosis in other spiders [13,27,38]. In *Mermessus trilobatus*, most (~70%) individuals were infected with the same strain of *Rickettsia* (R_4_), but we detected four other symbiont strains as well, frequently in conjunction with the common *Rickettsia* strain (Fig. 1b). In *G. foxi*, all specimens were infected with a previously uncharacterized bacterial OTU from the Order Rickettsiales (strain U1, Genbank Accession # MK529762) that had no close matches in the NCBI database (Fig. 1c). Nearly half of *G. foxi* specimens were additionally positive for a distinct *Wolbachia* strain that was not present in any of the other spider species (W_9_), and two *G. foxi* specimens also contained a distinct strain of *Rickettsiella* (RL_2_). In a subsequent diagnostic screen for *Rickettsiella*, we found 4/14 *G. foxi* to have *Rickettsiella*, whereas only a single specimen of *I. rugosa* (out of 51) was positive.

**Table 2.** Symbiotic microbiome profiles of individual spider specimens, based on the V4 segment of bacterial 16S rRNA

|  | Total | Symbiotypes of individual specimens |
| --- | --- | --- |
| *Agyneta unimaculata* | 5 | C<sub>2</sub>S<sub>1</sub>S<sub>2</sub>, R<sub>1</sub>S<sub>1</sub>S<sub>2</sub>, R<sub>1</sub>R<sub>8</sub>S<sub>3</sub>S<sub>4</sub>, 0 (×2)<sup>a</sup> |
| *Erigone autumnalis* | 7 | W<sub>10</sub> (×2), 0 (×5) |
| *Islandiana flaveola* | 2 | S<sub>1</sub>, 0 |
| *Mermessus* sp. 1 | 20 | W<sub>3</sub> (×16), W<sub>5</sub>, W<sub>10</sub>W<sub>12</sub>R<sub>1</sub>R<sub>7</sub>, 0 (×2) |
| *Mermessus tridentatus* | 1 | W<sub>3</sub>R<sub>6</sub><sup>b</sup> |
| *Tennesseellum formica* | 4 | W<sub>11</sub>, 0 (×3) |
| *Walckenaeria microspiralis* | 1 | C<sub>3</sub>W<sub>5</sub> |
| *Oxyopes salticus* | 9 | W<sub>7</sub> (×6), W<sub>8</sub>, 0 (×2) |

<sup>a</sup> Abbreviations: C = *Cardinium*, R = *Rickettsia*, S = *Spiroplasma*, W = *Wolbachia*, 0 = uninfected. Subscripts indicate strain type, commas separate symbiotypes, parenthetical numbers indicate number of specimens with the same symbiotype.
<sup>b</sup> Sanger sequencing of *fbpA* suggests a different and/or multiple *Wolbachia* strains.

**Fig. 1.**
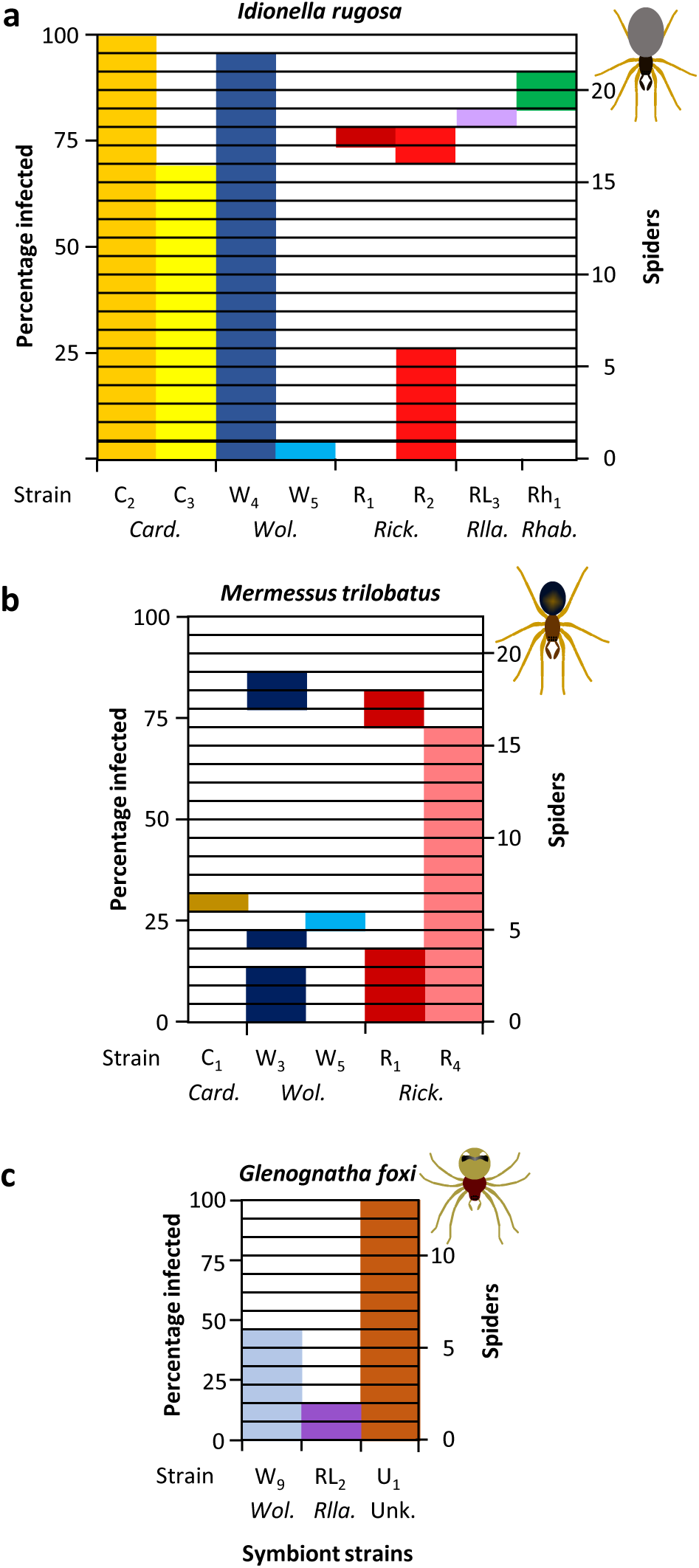
Symbiotypes of specimens of (a) *Idionella rugosa*, (b) *Mermessus trilobatus*, and (c) *Glenognatha foxi* based on Illumina MiSeq microbiome profiling of the V4 region of bacterial 16S rRNA. Each horizontal row corresponds to a single spider specimen: shaded boxes in each row indicate the specimen was positive for the corresponding symbiont for that column. Specimens with multiple shaded boxes were positive for a combination of symbiont strains. Symbiont abbreviations: C = *Cardinium*, W = *Wolbachia*, R = *Rickettsia*, RL = *Rickettsiella*, Rh = *Rhabdochlamydia*, U = an unknown bacteria in Rickettsiales. Different subscripts within a genus indicate OTUs that differ from one another by one or more basepairs

Other spider species were less variable in symbiotype (Table 2, [Online Resource 3]), although several individual specimens were positive for multiple symbiont strains. Four *Spiroplasma* OTUs were detected in pairs within specimens in *Agyneta unimaculata*, one of which was also in an *Islandiana flaveola* specimen [Online Resources 1, 3]. Subsequent diagnostic screens and Sanger sequencing of *A. unimaculata* and *I. flaveola* validated the presence of *Spiroplasma* in these specimens. In most specimens, endosymbiotic bacterial genera dominated the reads, constituting 70% of reads on average, but frequently reaching 98-99% [Online Resource 3]. Several additional OTUs that were not recognizably endosymbiotic were dominant in individual specimens [Online Resource 3]. Unlike OTU U1 in *G. foxi*, these were not widespread within or among host taxa, and thus we did not categorize them as potentially endosymbiotic at this time. In two unstarved *Oxyopes salticus* specimens we detected a strong bacterial signal of *Buchnera*, the obligate endosymbiont of aphids, which likely corresponded to gut contents [Online Resource 3]. Ten specimens, primarily in *Erigone autumnalis* and *Tennesseellum formica*, had no dominant OTUs, with all OTUs representing single digit or fractional percentages of the community. Finally, the amplification of the microbiomes of 17 specimens (primarily in *Agyneta* spp. *Florinda coccinea*, and *T. formica*), failed to generate sufficient high-throughput reads for analysis. Almost all of these samples (15) corresponded to samples that were diagnostically negative, reinforcing the interpretation that endosymbiont infections were not present in these specimens. For specimens that were included in multiple MiSeq sequencing runs, qualitative diagnoses of symbiont strain presence/absence were consistent between runs, as were quantitative proportions of reads that these OTUs represented per specimen [Online Resource 3].

In terms of strain diversity, *Wolbachia* strains exhibited much greater sequence divergence over the V4 region than either *Cardinium* or *Rickettsia*. In *Cardinium*, the three observed strains were identical in length and showed 99.2% pairwise similarity to one another over 251 basepairs, differing from one another by only one or two basepairs. Sanger sequencing slightly extended the sequenced region (by ~80 bases on the 3’ end), which yielded more variable sites, decreasing pairwise identity to 98.1% and reinforcing the distinction among the strains. *Rickettsia* also showed high similarity among strains over the V4 region, with 5 strains showing 98.4% pairwise identity, differing from each other by no more than 5 basepairs. However, longer Sanger sequencing of *Rickettsia* 16S that also included the V2 and V3 regions showed that two additional *Rickettsia* strains were present in the dataset, based on subtle differences between strains R_1_ and R_3_ (1 basepair) and R_1_ and R_6_ (2 basepairs). Strains R_1_, R_3_, and R_6_ were indistinguishable from one another based on the V4 region alone. For *Wolbachia*, we observed 9 different OTUs that had 96.7% average pairwise identity over 251 basepairs of 16S, differing by as much as 15bp from one another. We sequenced the *Wolbachia* surface protein (*wsp*) and/or fructose biphosphate adolase (*fbpA*) gene from 62 *Wolbachia*-positive specimens, and generally found strain type variants to be consistent with those based on 16S. However, in the single specimen from *Mermessus tridentatus*, *fbpA* sequence indicated a *Wolbachia* strain other than W_3_, even though the 16S sequence was identical to W_3_. When we inspected the chromatogram, weak double peaks suggested a multiple *Wolbachia* infection in *M. tridentatus*, but there was no sign of such in the high-throughput reads for this specimen. For three *Wolbachia* strains (W_3_, W_4_, and W_5_) we were able to sequence all 5 MLST genes used for *Wolbachia* strain typing [30]. Most sequenced alleles (13/15) were novel and not yet accessed to the PubMLST database. The most similar public alleles in the database indicated that our spider specimens contained *Wolbachia* strains from both Supergroups A and B [21], and that in five instances the closest or identical alleles came from *Wolbachia* strains that also inhabited spiders [24,30]. Overall, of the 27 endosymbiont strain types that we detected, most (21) were private to a single spider species [Online resources 1,3], but six were found in multiple spider species. Of the private strains, ten were each found in only a single specimen, five of which were unstarved, suggesting the potential for spurious positives due to symbiont-infected prey in the gut.

In general, starved specimens had a lower diversity of bacterial taxa than unstarved specimens, but the dominance of a few symbiotic taxa per specimen was unchanged. For both *I. rugosa* and *G. foxi*, there was no difference in the Shannon Diversity Index between starved and unstarved specimens, when rarified to a uniform read depth of 3000 reads (*I. rugosa*: H = 1.29 P = 0.25; *G. foxi*: H = 1.48, P = 0.22). For all specimens of both species, the bacterial community was dominated by only 1-5 OTUs, which constituted 75-99.9% of reads [Online Resources 3,4]. However, the number of OTUs represented in the remaining proportion of the sample was much lower in starved than unstarved specimens for both *I. rugosa* (median observed OTUs/3000 reads for starved = 12, unstarved = 60; H = 11.0 P<0.001) and *G. foxi* (median OTUs/3000 reads for starved = 11, unstarved = 39; H = 5.9, P = 0.007). Because unstarved specimens had many more rare taxa than starved specimens, PERMANOVA of the unweighted UniFrac metric showed significant community differences based on feeding status for both *I. rugosa* (pseudoF = 6.15, P < 0.001) and *G. foxi* (pseudoF = 2.75, P = 0.002). However, weighted Unifrac, which places more emphasis abundant taxa, did not differ based on feeding status for either species (*I. rugosa*: pseudoF = 1.19, P = 0.20; *G. foxi*: pseudoF = 0.68, P=0.58).

## Discussion

Spiders have diverse endosymbiotic microbiomes. In a survey of 267 agricultural spiders representing 14 species, we found 27 different endosymbiont strain types, representing 7 bacterial genera. The majority of spider species had individuals that tested positive for endosymbionts, and several species had individuals that tested positive for two or more symbiont strains simultaneously. The most symbiont-rich spider species, *Idionella rugosa*, had 8 different strain types representing 5 endosymbiotic bacterial genera. This rivals or exceeds the endosymbiont diversity observed in insects [14,39], particularly given the limited geographic, temporal, and habitat diversity of spiders sampled in the current study. Most symbiont strains were widespread within and characteristic to a particular spider species, but there was substantial variation in symbiont profiles among specimens, suggesting the potential for both interactions among symbionts within hosts, and dynamic interactions of differentially infected individuals within populations.

Spiders also have novel endosymbiont taxa that have not been previously described. We found that all specimens of *G. foxi* were infected with the same OTU that falls within the Rickettsiales, the same order as *Wolbachia* and *Rickettsia*, but not within the families of either of these well-known endosymbiotic bacterial genera. The signal from this novel Rickettsiales was very strong, constituting the vast majority of reads in the microbiome profiles of most *G. foxi* specimens [Online Resource 3]. These specimens were collected over multiple years and throughout the season, suggesting a chronic and persistent infection in the population, not a brief pathogenic outbreak. Further work would be necessary to determine transmission methodology and fitness consequences of infection, but a preliminary hypothesis of a novel symbiotic clade seems reasonably supported. Another non-*Wolbachia* Rickettsiales was recently found to induce cytoplasmic incompatibility in a beetle [40], indicating that our understanding of the breadth of endosymbiotic associations within the Rickettsiales is still expanding. We additionally observed a strain of *Rhabdochlamydia*, and two different strains of *Rickettsiella* within spider specimens. Both of these genera have previously been reported in linyphiid spiders [13,27, 38], and the latter genus has been associated with both mutualistic and pathogenic function in other arthropods [41–43].

In general, the function of these endosymbiotic taxa in spiders remain to be elucidated, although many are likely to manipulate spider reproduction. In *Mermessus fradeorum*, several endosymbiont strains are present, causing both cytoplasmic incompatibility and feminization [26]. Other studies have similarly associated spider endosymbionts with reproductive manipulation [25,44] or found increased prevalence of endosymbionts in female over male specimens [22]. However, yet other studies have found no obvious reproductive manipulations associated with spider endosymbionts [45]. As with endosymbionts in insects, it can be expected that increased attention to the spider microbiome will reveal an expanded repertoire of phenotypic effects that these symbionts can invoke [46]. Already, there are suggestions that spider endosymbionts may induce novel reproductive and dispersal effects in their hosts [47,48].

In the present study we included both starved and unstarved specimens, to gain insight into the potential utility of the latter. Starved specimens had fewer microbial taxa than unstarved, but it appears that it is mostly low-representation taxa that are removed from the microbiome by starvation. With respect to dominant bacterial taxa in the samples, starved and unstarved specimens were quite similar, typically characterized by the same few symbiotic taxa. Thus, we would have likely come to the same major conclusions had we worked with only unstarved specimens. We conclude that unstarved specimens would likely be sufficient for studies that are focused on the distribution of characteristic endosymbionts within a host species, and that molecular samples collected for other ecological or phylogenetic studies [e.g.,28,29] could provide a rich repository for investigating symbiont infection in populations over time and space.

However, it was useful to also have starved specimens in the dataset. When starved and unstarved specimens had concordant endosymbiont strain types, this served as confirmation that the positive diagnoses from unstarved specimens were genuine. Such confirmation was particularly necessary for symbionts that occurred relatively infrequently in a host population (e.g. *Rhabdochlamydia* in starved *I. rugosa*), which otherwise might be rightly viewed with skepticism. The presence of the aphid obligate endosymbiont *Buchnera* in some of our unstarved specimens demonstrates the very real potential for false positives via gut contents. It is worth remembering that spiders are frequent cannibals and intraguild predators [49,50], which could result in characteristic symbionts in their digestive systems due to recent consumption of infected conspecifics. Thus, starved specimens, or specimens reared on a controlled symbiont-free diet, remain the gold standard for symbiont characterization. Further work is needed to understand the tenure of consumed symbionts in the spider digestive system, as well as the potential for horizontal transfer of symbionts from prey to predator [51].

It was also beneficial to include a combination of diagnostic PCR, Sanger sequencing, and high-throughput sequencing in our survey. Diagnostic PCR, even in combination with Sanger sequencing, underestimated endosymbiont diversity, both by missing unexpected bacterial taxa (e.g., OTU U1 and *Rickettsiella* in *G. foxi*), and by failing to resolve infections by multiple congeneric strain types within the same host specimen (e.g., the two *Cardinium* strains in *I. rugosa*). However, the high-throughput data also had limitations. In particular, index swapping and ultra-sensitivity of the methodology had a tendency to result in false positives, unless data was corrected [36]. Furthermore, the short amplification region failed to fully resolve some strain types that were evident from Sanger sequencing of longer segments of 16S or alternate genes. The V4 region, and 16S as a whole, has previously been shown to be problematic for some taxonomic groups that are invariant over this region (e.g. many genera in the Enterobacteriaceae), and also to over-represent taxa that have multiple non-identical 16S copies within their genome [52, 53]. Using both approaches, we were able to capitalize on their complementary strengths. Together, they revealed a tremendous diversity of endosymbionts within common agricultural spiders, and suggest that this will be a fruitful system for further investigations of interactions among endosymbionts.

## Supporting information

Online Resource 1

Online Resource 2

Online Resource 3

Online Resource 4

## Acknowledgements

We thank J. Harwood and J. Dryer for providing and collecting specimens. This research work was supported by grants from the Kentucky Science and Engineering Foundation as per Grant/Award Agreements 148-502-10-261 and 148-502-16-377 with the Kentucky Science and Technology Corporation, and the National Institute of Food and Agriculture, U.S. Department of Agriculture (Hatch No. 0224651).

## Electronic supplementary material

Online Resource 1: Individual spider specimens, diagnostic results, and sequences. Tab1 includes metadata, diagnostic results, COI haplotypes and bacterial strain types associated with each specimen. Tab 2 provides the Genbank accession number and sequences associated with each COI haplotype and bacterial strain type.

Online Resource 2: Diagnostic PCR primers.

Online Resource 3: High throughput read distribution per sample.

Online Resource 4: **Figs S1 & S2** Microbiome profile of starved versus unstarved specimens of (**Fig. S1**) *Idionella rugosa* and (**Fig. S2**) *Glenognatha foxi*. Each profile was generated from a rarified sample of 3000 Illumina MiSeq reads of the V4 region of bacterial 16S rRNA. The 7-8 most common genera are presented in the key for each figure; in **Fig. S1**, *Cardinium, Wolbachia* and *Rickettsia* each encompass two distinct strain type OTUs. In **Fig. S2**, many of these OTUs could only be placed at the family or order level.

