## Supplementary material for "Endosymbiotic bacteria are prevalent and diverse in agricultural spiders": Online Resource 2

Online Resource 2. Primers used for diagnostic PCR.

| Target | Target gene | Primer name | Primer sequence 5' to 3' | Expected product size (bases) | Annealing temp | References |
|---|---|---|---|---|---|---|
| *Cardinium* | 16S | CHF<br>CHR | TACTGTAAGAATAAGCACCGGC<br>GTGGATCACTTAACGCTTTCG | ~300 | 56°C | Zchori-Fein and Perlman 2004 |
| *Rickettsia* | 16S | 16saiF<br>Rick16sR | AGAGTTTGATCMTGGCTCAG<br>CATCCATCAGCGATAAATCTTTC | ~200 | 60°C | Fukatsu and Nikoh 1998<br>Fukatsu et al. 2001 |
| | 16S | RicklongF<br>RicklongR | ACGTGGGAATCTACCCATCA<br>TAGCCTAGATGACCGCCTTC | ~500 | 60°C | Curry et al. 2015 |
| *Rickettsiella* | 16S | RLA16s F1<br>RLA16s R1 | CAGTAAARRTTTCGGYCTTTAYGGG<br>CAAACCTAGTCAACCACCTACACG | ~530 | 56°C | Duron et al. 2016 |
| *Spiroplasma*[1] | 16S | 16saiF<br>TKsSSpr | AGAGTTTGATCMTGGCTCAG<br>TAGCCGTGGCTTTCTGGTAA | ~350 | 60°C | Fukatsu et al. 2001 |
| *Wolbachia* | fbpA | fbpAF1<br>fbpAR1 | GCTGCTCCRCTTGGYWTGAT<br>CCRCCAGARAAAAYYACTATTC | ~500 | 59°C | Baldo et al. 2006 |
| | wsp | wspF1<br>wspR1 | GTCCAATARSTGATGARGAAAC<br>CYGCACCAAYAGYRCTRTAAA | ~500 | 59°C | Baldo et al. 2006 |
| spider | COI | lco1490<br>hco2198 | GGTCAACAAATCATAAAGATATTGG<br>TAAACTTCAGGGTGACCAAAAAATCA | ~650 | 53°C | Folmer et al. 1994 |

[1]Prone to amplifying other bacterial genera in addition to *Spiroplasma*. Sanger sequencing for validation is requisite.
