## Supplementary material for "Endosymbiotic bacteria are prevalent and diverse in agricultural spiders": Online Resource 4

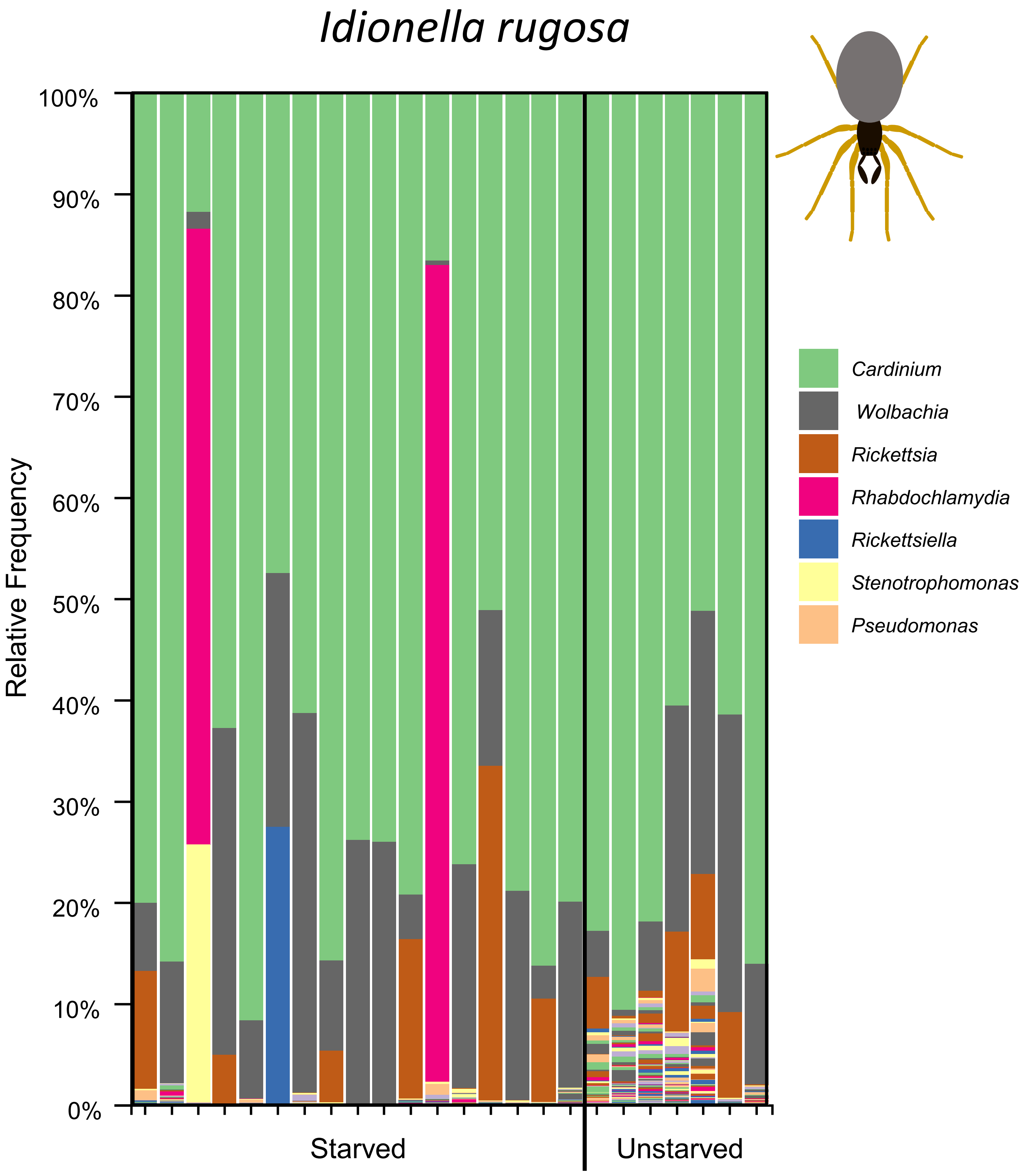


Figure S1. Microbiome profile of starved versus unstarved specimens of *Idionella rugosa.* Each profile was generated from a rarified sample of 3000 Illumina miseq reads of the V4 region of bacterial 16S sequence. The 7 most common genera are presented in the key; *Cardinium, Wolbachia* and Rickettsia each encompass two distinct strain type OTUs.

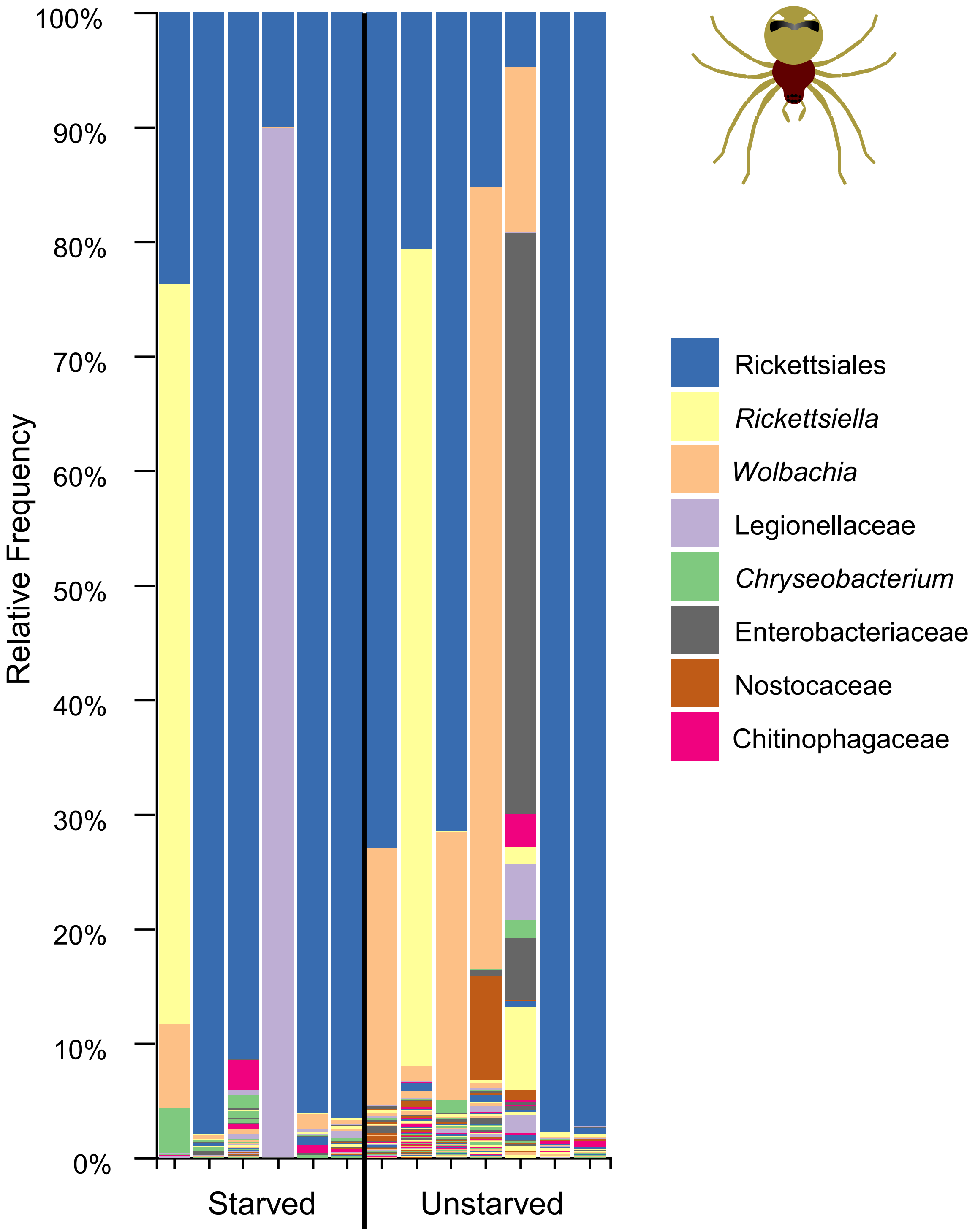


Figure S2. Microbiome profile of starved versus unstarved specimens of *Glenognatha foxi.* Each profile was generated from a rarified sample of 3000 Illumina miseq reads of the V4 region of bacterial 16S sequence. The 8 most common genera are presented in the key, many of which could only be placed at the family or order level.
